# Transcript-specific determinants of pre-mRNA splicing revealed through *in vivo* kinetic analyses of the 1^st^ and 2^nd^ chemical steps

**DOI:** 10.1101/2020.10.13.338152

**Authors:** Michael A. Gildea, Zachary W. Dwyer, Jeffrey A. Pleiss

## Abstract

Understanding how the spliceosome processes its composite of substrates through the two chemical steps required for mRNA production will be essential to deciphering splicing regulation and mis-regulation. Here we measure the *in vivo* rates of these steps across the genome by coupling metabolic RNA labeling, targeted sequencing, and first order kinetic modeling. We reveal a wide variety of rates by which introns are removed, that splice site sequences are primary determinants of 1^st^ step rates, and that the 2^nd^ step is generally faster than the 1^st^ step. We show that ribosomal protein genes (RPGs) are spliced faster than non-RPGs at each step, and that RPGs share evolutionarily conserved cis-features which facilitate their splicing. A genetic variant defective at the 1^st^ step shows the expected defect in the 1^st^ step, but an unexpected change in the 2^nd^ step which suggests how co-transcriptional splicing functions as an important determinant of splicing rates.

## Introduction

In the time since the discovery of pre-messenger RNA (pre-mRNA) splicing, our understanding of its importance in regulating gene expression has grown dramatically. This is perhaps best exemplified by the ever-increasing number of human diseases that are found to be associated with mutations in the splicing pathway (Montes et al., 2019, Scotti and Swanson, 2016). Critical to our understanding of these diseases is a comprehensive knowledge of the mechanisms by which the spliceosome processes its diverse set of substrates: for many such diseases, the fundamental question of whether the disease-state results from a defect in the splicing of the global complement of transcripts, or in the splicing of a specific but critical subset of transcripts, remains unresolved. Importantly, many of these details remain unclear due to the difficult problem the complexity of splicing regulation presents. Pre-mRNA splicing requires that the spliceosome, which catalyzes intron removal, accurately defines, assembles upon, and activates the appropriate splice site sequences in the background of a sea of non-cognate, cryptic sites (Lee and Rio, 2015, Wahl et al., 2009). Moreover, the spliceosome must catalyze the reaction in a timely manner, striking a balance between fidelity and speed to achieve maximal efficiency (Semlow and Staley, 2012). An understanding of the relative speeds or rates with which the complement of spliceosomal substrates are processed will allow for elucidation of substrate specific features that enable their regulation.

The spliceosome removes introns by catalyzing two step-wise transesterification reactions. Decades of research has established that cis sequence elements within introns and transcripts influence splicing outcomes. Notably, the 5’ splice site (5’SS), 3’ splice site (3’SS), and branch point (BP) sequences define introns by directly base pairing to partially complimentary sequences within the spliceosomal snRNAs, and as a result couple cis-elements within an intron with splice site selection (Wahl et al., 2009). However, the influence of these sequences, among other cis-elements, on the rate with which an intron is processed through each chemical step of splicing *in vivo* is poorly understood. Though many advancements have been made in our understanding of splicing and the spliceosome through detailed structural and functional studies, methodological limitations have made it difficult to deconvolute the influence of cis-features on the rates of the 1^st^ and 2^nd^ steps of splicing *in vivo* (Fica and Nagai, 2017, Shi, 2017, Lee and Rio, 2015, Wahl et al., 2009, Mayerle and Guthrie, 2017). For example, RNA-seq is commonly used to assess the impact of specific cis and/or trans-acting factors on splicing efficiency by comparing changes in reads representing unspliced and spliced transcripts. Yet, as commonly employed this approach is intrinsically limited by its reliance on steady-state measurements. While splicing rate and fidelity influence the steady-state abundances of unspliced and spliced species, so too do other processes such as degradation of the mature species, making it impossible to deconvolute changes in splicing rate from other degradation rates using steady-state measurements alone (Wachutka and Gagneur, 2017). By contrast, many investigators historically used cellular extracts to measure the kinetics of *in vitro* splicing of reporter constructs, avoiding the pitfalls of steady state analyses (Mayerle and Guthrie, 2017, Hicks et al., 2005). But while such studies have provided important insights into the splicing process, it remains unknown how the results from these studies, where fully formed transcripts are presented to an unfractionated mixture of cellular components, relate to *in vivo* conditions across the global complement of substrates.

In addition to the role of *cis*-regulatory elements on splicing efficiency, it is well established that other pre-mRNA processing events are functionally coupled to splicing (Herzel et al., 2017, Naftelberg et al., 2015, Bentley, 2014). Current data overwhelmingly support a model wherein splicing is temporally, spatially, and functionally connected with transcription (Herzel et al., 2017, Naftelberg et al., 2015, Bentley, 2014). Indeed, *in vivo* studies have demonstrated the capacity of transcription to influence splicing outcomes (Naftelberg et al., 2015, Bentley, 2014). Nevertheless, estimates of when splicing completes in relation to transcription range widely. Whereas the seminal studies which established the co-transcriptional nature of splicing showed that assembly of the spliceosome occurs in a largely co-transcriptional fashion, these studies nevertheless concluded that the chemical steps generally complete post-transcriptionally (Lacadie et al., 2006, Tardiff et al., 2006, Moore et al., 2006). More recently however, studies using a variety of methods have yielded widely disparate estimates of when splicing completes with respect to the position of RNA polymerase. For example, whereas work from the Neugebauer group suggested that the chemistry of splicing may complete almost concurrently with production of the 3’SS (Oesterreich et al., 2016, Reimer et al., 2020), work from the Churchman group using a similar method suggested that the chemistry of splicing generally completes after kilobases of RNA have been transcribed (Drexler et al., 2020). A clearer picture of the functional coupling of splicing and transcription as well as the influence of splicing on gene expression would be bolstered by robust measurements of splicing kinetics at individual step resolution.

Recently, the use of metabolic RNA labeling to estimate RNA processing rates *in vivo* has gained popularity and greatly expanded our knowledge of pre-mRNA processing kinetics in a variety of organisms (Duffy et al., 2019). This method enables the time-resolved isolation of nascent RNA and utilizes kinetic modeling of approach to equilibrium curves to estimate the rates of production and degradation of different RNA isoforms. This approach has been applied to measure splicing rates in a variety of organisms, generally using standard RNA-seq methods (Barrass et al., 2015, Wachutka et al., 2019, Pai et al., 2017, Eser et al., 2016, Rabani et al., 2014, Windhager et al., 2012). Despite these advances, the uniformity of RNA-seq read coverage across transcripts produces a low abundance of splicing informative reads per splicing event, resulting in poor quantification of many splicing events and thus their splicing rates. To improve upon these limitations, our lab has recently developed Multiplexed Primer Extension sequencing, or MPE-seq, a targeted RNA-seq method that provides a significant enrichment for splicing informative reads in RNA-seq data sets and enables differentiation of completely unspliced, lariat intermediate, and spliced isoforms genome-wide (Xu et al., 2019, Gildea et al., 2019). Here, we report on our measurements of the 1^st^ and 2^nd^ step splicing rates for the genome-wide compliment of substrates in the budding yeast *Saccharomyces cerevisiae* that were generated by combining rapid metabolic RNA labeling, MPE-seq, and modeling of approach to steady state kinetics.

## Results

To better understand the relative efficiencies by which the spliceosome processes its composite of substrates, we designed a strategy to determine the genome-wide rates of the 1^st^ and 2^nd^ steps of splicing. As outlined in Figure 1, our approach incorporated three principle components, briefly described here and more fully described in the Methods section. First, we employed a rapid metabolic labelling approach using 4-thiouracil (4tu) to isolate nascent RNA at time intervals ranging from seconds to minutes. Second, we used multiplexed primer extension sequencing (MPE-seq) to quantify splicing intermediates and track their processing through time. Our lab has previously demonstrated two important properties of MPE-seq that are essential to this work: a dramatic increase in the sensitivity for detecting splicing-informative reads, enabling high precision measurements of their relative abundances; and the capacity to distinguish between unspliced, lariat intermediate, and spliced isoforms of a given transcript.

**Figure 1.**
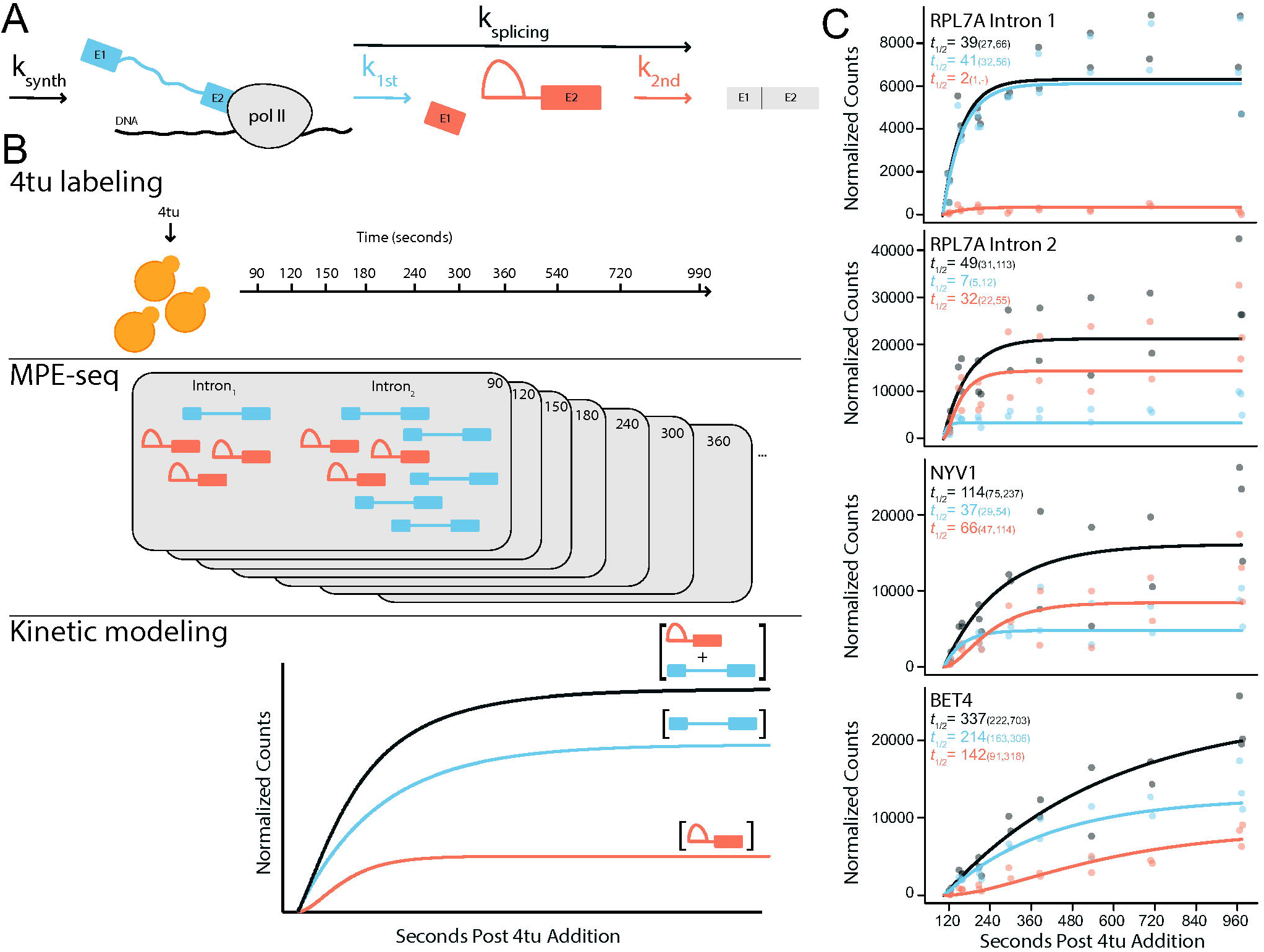
Determining the rates of 1^st^ and 2^nd^ Chemical Step of pre-mRNA Splicing. (A) A schematic of the rates measured in this study. (B) Experimental workflow. 4-thiouracil (4tu) is introduced to actively growing yeast cells followed by sample collection at specified time points. Nascent 4tu-labeled RNA is purified from each sample and MPE-seq libraries are prepared. The pre 1^st^ step and lariat intermediate isoforms are quantified genome wide, and kinetic models are fit to the approach to equilibrium curves to estimate the total (grey), 1^st^ (blue), and 2^nd^ (orange) step rates of pre-mRNA splicing. (C) Normalized counts of total unspliced (grey), pre 1^st^ step (blue), lariat intermediate (orange) versus time in seconds after addition of 4tu for RPL7A intron 1, RPL7A intron 2, NYV1, and BET4. Solid lines are model fits and corresponding half-lives are provided with 90% confidence intervals in parenthesis.

Finally, these data were fit to first order kinetic models to estimate the half-lives of the 1^st^ and 2^nd^ steps of splicing genome-wide: the high resolution afforded by MPE-seq allowed us to robustly measure the change in abundance of these three splicing isoforms for each intron in the genome through time, enabling generation of high-quality kinetic models for these rates.

### Measuring genome-wide splicing rates in vivo

Our method was first employed on exponentially growing wild type (BY4741) *S. cerevisiae* cells grown in synthetic medium at 22°C. Time points after 4tu addition were chosen at intervals as short as 30 seconds in order to maximize the number of splicing events for which sufficient data could be generated within the approach to steady state curves to enable high confidence model fits. We note that 22°C is a standard temperature for yeast growth in the wild, and while many laboratory studies use 30°C to optimize growth rate, our initial experiments suggested that we could better model the kinetic data generated at 22°C (see also Methods and Discussion for further consideration). To measure the absolute abundance of each RNA species in each sample, a fixed amount of an exogenous pool of *in vitro* transcribed, 4-thiouridine (4su) containing RNAs was added to each sample as a spike-in control prior to RNA purification. As expected, the abundance of 4su-containing RNAs isolated from our total yeast RNA pools increased over time (Figure S1A), consistent with increasing incorporation of 4su in nascent RNAs. MPE-seq libraries were prepared from the 4su-containing RNA purified from each sample, and the libraries were sequenced using paired-end sequencing on an Illumina NextSeq platform. Roughly 10 million total reads were obtained for each of three replicate samples of each time point (Table S1), equivalent to ∼1 billion reads of standard RNAseq per individual replicate sample (Gildea et al., 2019). Sequencing reads were aligned and those representing the pre 1^st^ step, lariat intermediate, and spliced isoforms of each intron-containing gene were quantified. To account for variations in processing efficiency between time point and replicate samples, the counts for each species were normalized to the read counts corresponding to the spike-in control RNAs. Normalized reads for each species in the samples collected prior to addition of 4tu to the media were used to estimate and correct for the background signal for each RNA species in each sample. Consistent with the isolation of nascent RNA, the fraction of reads corresponding to unspliced RNA (pre 1^st^ step + lariat intermediate) relative to spliced RNA per intron was highest in the early time points and decreased over time (Figure S1B).

To determine the rates of each chemical step in the splicing pathway, we used first-order kinetic models to fit the approach to equilibrium curves generated for each of the spliced isoforms.

Simply stated, these models describe how the rate of increase in abundance of a specific RNA species over time becomes balanced by decay due to splicing and/or degradation (see methods section for a detailed description of the models and model fitting procedures). In order to determine the overall splicing rates, as well as the rates of the 1^st^ and 2^nd^ chemical steps for each splicing event, we established an initial set of definitions. First, the synthesis rate for each intron was defined as the number of complete introns synthesized per unit time. Second, the overall splicing rate was defined as the time from completion of synthesis to completion of the 2^nd^ chemical step of splicing. Third, the rate of the 1^st^ chemical step was defined as the time from completion of synthesis to completion of the 1^st^ chemical step. And finally, the rate of the 2^nd^ chemical step was defined as the time from completion of the 1^st^ chemical step to completion of the 2^nd^ chemical step. In addition to these guiding definitions, a constant time offset was included due to the delay in the time between the addition of 4-thiouracil (4tu) to the growth medium and the steady state availability of 4-thiouridine (4su) to the transcription machinery in the nucleus. Using this approach, the abundance of total unspliced counts were modeled to estimate the total rate of splicing, akin to other RNA-seq based kinetics studies (Eser et al., 2016). The abundance of pre 1^st^ step and lariat intermediate counts were similarly modeled to estimate the rates of the synthesis, 1^st^ and 2^nd^ steps of splicing. Splicing rates are represented as half-lives throughout this manuscript. High confidence synthesis, total, 1^st^, and 2^nd^ step rate estimates were obtained for the majority of introns in the genome (Figure 1B and Table S2). Confidence intervals for parameter estimates were used to evaluate model fits (Table S2).

### Genome-wide rates reveal a wide variation in splicing efficiency, and that the 2^nd^ step is generally faster than the 1^st^ step

We first assessed the global distributions of the half-lives for the total, 1^st^ step, and 2^nd^ step of pre-mRNA splicing. A median half-life of 135, 83, and 41 seconds was estimated for each of these steps, respectively (Figure 2A). Remarkably, these data revealed a >30-fold variation in individual splicing rates across the genome (Figure 2A). Additionally, a large variation in relative 1^st^ and 2^nd^ step rates were apparent between individual introns, including in some cases even between two introns housed within the same transcript. For example, both introns in the Rpl7A pre-mRNA were excised with similar overall splicing rates, however, that overall rate was achieved through contrasting 1^st^ and 2^nd^ step rates (Figure 1C), suggesting that the spliceosome and its substrates may have evolved similar total rates of pre-mRNA splicing through contrasting means. Nevertheless, when considering all spliceosomal substrates, no significant correlation was observed between the measured 1^st^ and 2^nd^ step rates (Figure S2).

**Figure 2.**
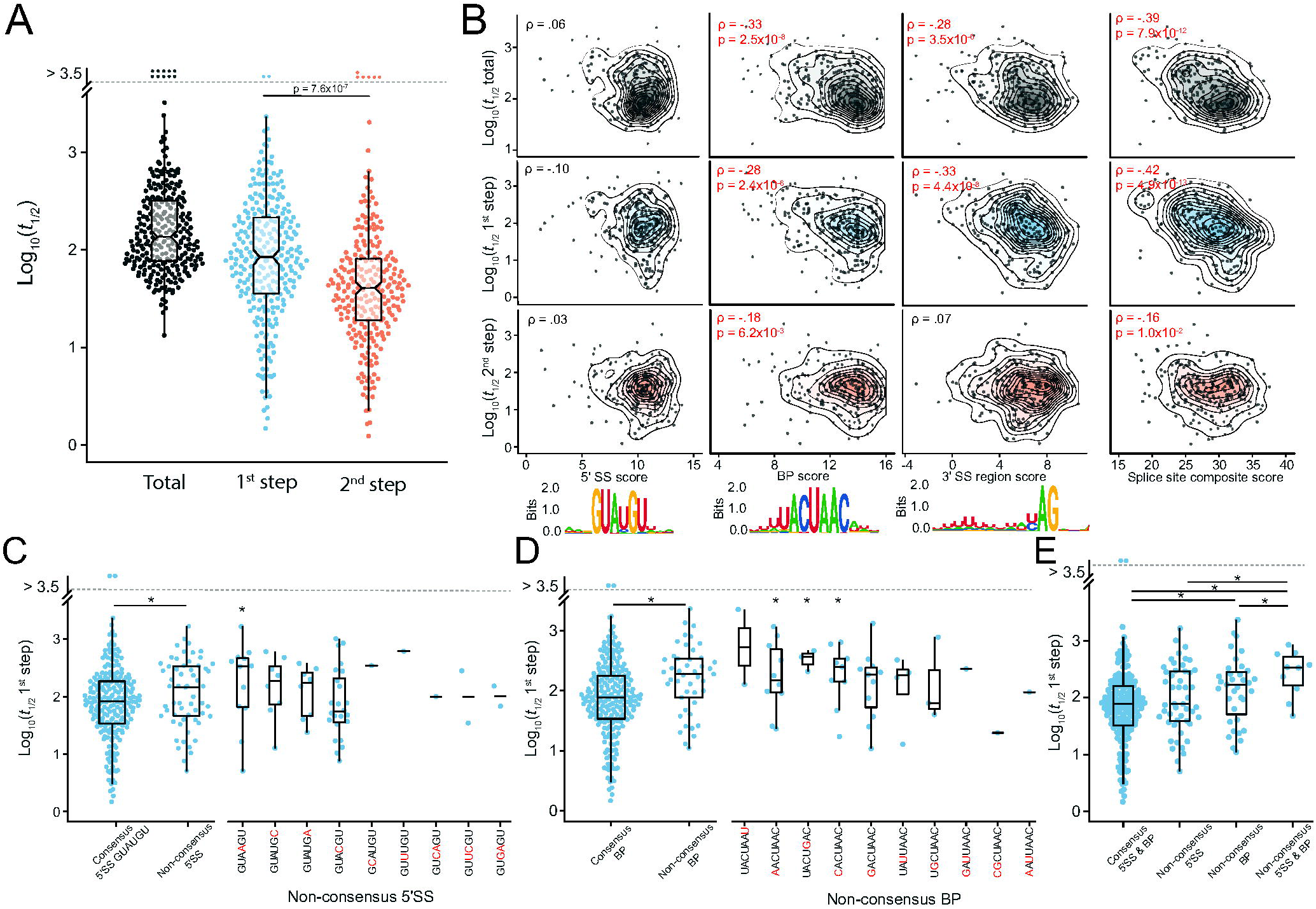
Splice Site Sequences are Major Determinants of the Rate of the 1^st^ Step of pre-mRNA Splicing. (A) Distributions of the half-lives for the total (n = 278), 1^st^ step (n = 272), and 2^nd^ step (n = 241) of pre-mRNA splicing. The 2^nd^ step on average is significantly faster than the 1^st^ step (B) Comparison of 5′SS, BP, 3′SS, and composite splice site scores with total, 1^st^, and 2^nd^ step rates. Web logos for 5′SS, BP, and 3′SS are included. Spearman correlation coefficients (ρ) and associated p-values (p) are included. Red text and highlighted axes indicate significant correlations (p < 0.05). (C) Comparison of 1^st^ step splicing rates between introns with consensus (n = 213) and non-consensus (n = 59) 5′SS sequences. Non-consensus 5′SS sequences are further partitioned by sequence. (D) Comparison of 1^st^ step splicing rates between introns with consensus (n = 227) and non-consensus (n = 45) BP sequences. Non-consensus BP sequences are further partitioned by sequence. (E) Comparison of 1^st^ step splicing rates between introns with consensus 5′SS and BP sequences, introns with non-consensus 5′SS and consensus BP sequences, introns with consensus 5′SS and non-consensus BP sequences, and introns with non-consensus 5′SS and non-consensus BP sequences. Statistical significance in panels C-E was calculated with one-sided Mann-Whitney signed-rank test. Statistical significance for panel A was calculated with one-sided Wilcoxon signed-rank test. *p<0.05.

### Cis-transcript features contribute to splicing kinetics

In order to understand what drives the observed variation in splicing rates, we investigated the influence of cis transcript features with overall and individual step splicing rates. We first investigated the influence of conserved cis-elements within introns that are recognized by components of the spliceosome and known to be crucial for efficient splicing, including the 5’ splice site (5’SS), branch point (BP), and 3’ splice site (3’SS) sequences. To investigate the influence of these sequences we calculated position weight matrix (PWM) based scores for the 5’SS, BP, and 3’SS regions and compared them to the measured half-lives for the total, 1^st^, and 2^nd^ steps (Table S3). We observed a significant correlation between the scores of the BP and 3’SS region sequences and the half-lives of both the total and 1^st^ steps of splicing (Figure 2B). By contrast, only a mild correlation was observed between the BP scores and the half-lives of the 2^nd^ step. Together, these data suggest that splice site strength is primarily a determinant of 1^st^ step rates. The composite of all three splice site scores showed an increased correlation with the half-lives of the total and 1^st^ steps, indicating that these sequences act to either additively or synergistically influence the 1^st^ step splicing rate (Figure 2B).

Surprisingly, a significant correlation was not observed between the 5’SS scores and the half-lives of the 1^st^ step (Figure 2B). However, we note that *S. cerevisiae* introns are defined by highly conserved splice site sequences, and over 75% of introns contain a GUAUGU sequence at their 5’SS. Indeed, we saw that those introns with the consensus 5’SS were spliced significantly faster than those with non-consensus 5’SS sequences (Figure 2C), suggesting that the lack of correlation observed for the 5’SS scores likely reflects an absence of resolving power in the PWM scores. Interestingly, by further comparing individual non-consensus 5’SS and 1^st^ step rates, we found that a substitution at the 4^th^ position of the 5’SS from a U to an A, which results in perfect complementarity to the U1 snRNA, resulted in the longest 1^st^ step half-lives of the non-consensus 5’SS for which there exists a sufficient sample size (Figure 2C). By contrast, a substitution at the 4^th^ position from a U to a C, which is the most common non-consensus 5’SS and retains a mis-match with the U1 snRNA, showed no significant difference in 1^st^ step half-life from the consensus 5’SS (Figure 2C).

A similar analysis was performed for the BP consensus (UACUAAC) sequence, and as expected, non-consensus BP sequences resulted in a significantly longer 1^st^ step half-life (Figure 2D). We did not observe a significant difference in 1^st^ step half-lives between individual non-consensus BP sequences (Figure 2D), however small sample sizes may have obfuscated meaningful differences. Introns with both non-consensus 5’SS and BP sequences were spliced significantly slower than all other introns and the magnitude of the difference was greater than for introns with only one non-consensus sequence (Figure 2E). The difference between introns with a non-consensus 5’SS and a consensus BP and those with both consensus 5’SS and BP was not statistically significant, likely because of differences in individual non-consensus 5’SS sequences (Figures 2C and 2E).

Finally, we assessed the role of the 3’SS (YAG), as well as a short (six nucleotide) poly-U tract just upstream of the 3’SS. PWM scores were generated using bases -6 through -11 from the 3’SS for the poly-U tract, while the YAG plus 2 bases upstream and 3 bases downstream were used in determining the 3’SS score (Figure S3A). The strength of both the poly-U tract and 3’SS showed a significant correlation with the half-lives of the 1^st^ step of splicing but not the 2^nd^ (Figure S3B).

### Ribosomal protein genes are spliced faster at both steps

The set of intron-containing genes in *S. cerevisiae* can be partitioned into 2 functional classes: ribosomal protein genes (RPGs) and non-ribosomal protein genes (nRPGs). Of the roughly 280 annotated introns in *S. cerevisiae*, more than one third are found within RPGs. RPGs are highly expressed under normal conditions, and in response to environmental changes or stress they have been shown to be coordinately regulated both at the level of transcription and splicing(Parenteau et al., 2011, Pleiss et al., 2007a, Parenteau et al., 2019, Gasch et al., 2000, Causton et al., 2001, Reja et al., 2015). Consistent with previous studies, we find that the total rate of splicing is fast for the RPGs relative to the nRPGs, with median half-lives of 84 and 218 seconds, respectively (Figure 3A) (Barrass et al., 2015). Additionally, a significantly higher synthesis rate was observed for RPGs compared to nRPGs (Figure 3B), consistent with their high level of expression. Whereas previous studies lacked the resolution to investigate the two individual steps of splicing, here we saw that splicing of RPGs was significantly faster at both the 1^st^ and the 2^nd^ steps of splicing (Figure 3A). Remarkably, the half-lives of the 1^st^ and 2^nd^ steps showed a significant negative correlation with one another within the RPG introns but not in the nRPGs introns, suggesting that RPGs may have evolved mechanisms to homogenize total splicing rates through optimization of different steps (Figure S4A). Consistent with their coordinated regulation, the variation in total splicing rates within the RPG class is considerably smaller when compared to the nRPGs.

**Figure 3.**
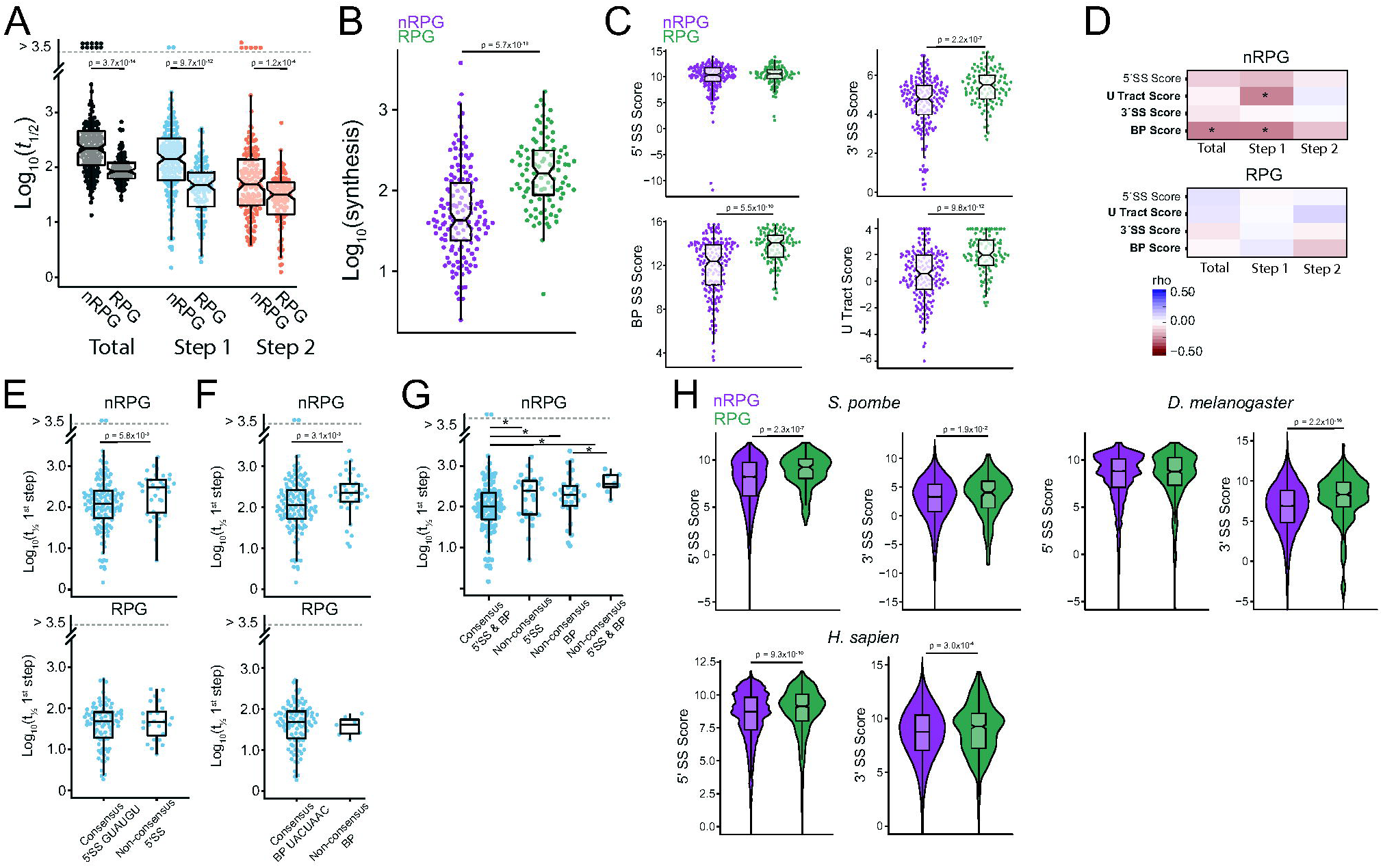
Both the Properties and the Splicing of RPG Introns is Distinct from nRPG Introns. (A) Distributions of the half-lives for the total (RPG n = 105, nRPG n = 173), 1^st^ step (RPG n = 105, nRPG n = 167), and 2^nd^ step (RPG n = 99, nRPG n = 142) of pre-mRNA splicing. (B) Synthesis rates for RPGs (n = 105) and nRPGs (n =173). (C) The PWM scores for 5′SS, BP, 3′SS, and U-tract regions compared between RPGs and nRPGs. (D) Spearman correlations between 5′SS, BP, 3′SS, and U-tract scores and half-lives for the total, 1^st^ step, and 2^nd^ step of pre-mRNA splicing for RPGs and nRPGs. (E) Comparison of the half-lives for the 1^st^ step of splicing between introns with consensus (RPG n = 81, nRPG n = 137) and non-consensus (RPG n = 24, nRPG n = 40) 5′SS sequences. (F) Comparison of half-lives for the 1^st^ step of splicing between introns with consensus (RPG n = 98, nRPG n = 135) and non-consensus (RPG n = 7, nRPG n = 42) BP sequences. (G) Among nRPG introns, a comparison of the half-lives of the 1^st^ step of splicing between those with consensus 5′SS and BP sequences, those with non-consensus 5′SS but consensus BP sequences, those with consensus 5′SS but non-consensus BP sequences, and those with non-consensus 5′SS and BP sequences. (H) The PWM scores distributions for 5′SS and 3′SS among RPG and nRPG introns in *S. pombe* (RPG n = 134, nRPG n = 5160), *H. sapiens* (RPG n = 801, nRPG = 423110), and *D. melanogaster* (RPG n = 499, nRPG = 149895). Statistical significance in panels A-C, and D-H was calculated with one-sided Mann-Whitney signed-rank test. *p<0.05.

To better understand what drives the relative efficiency of the 1^st^ and 2^nd^ steps of RPG splicing, and building off of previous observations that RPG introns have distinct features compared to nRPG introns (Spingola et al., 1999, Parker and Patterson, 1987), we asked how cis transcript features differed between RPGs and nRPGs. Notably, the RPGs as a class contained significantly stronger BP, 3’SS, and poly-U tract scores when compared to nRPGs, whereas no significant difference was observed in 5’SS scores (Figures 3C and S4B). We then asked whether these features were correlated with the observed splicing rates for the RPGs and nRPGs as independent groups. Interestingly, within the nRPGs, which as a class have lower scoring splice site sequences, we observed significant correlations between BP scores and the half-lives of the total and 1^st^ step, and U-tract scores with the half-lives of the 1^st^ step (Figure 3D). By contrast, no correlations were apparent between the rates of RPG splicing and any of these features. Moreover, within the nRPG group, introns with non-consensus 5’SS were spliced significantly slower compared to those with consensus sequences, but somewhat surprisingly no such difference was seen within the RPGs (Figure 3E); here we note, however, that the most common non-consensus 5’SS in RPGs is a variant harboring a C at the 4^th^ position (15 of the 24 variants) which also showed little impact on 1^st^ step half-lives in nRPGs. Similarly, introns with non-consensus BP sequences were also spliced significantly slower than those with consensus sequences within the nRPG class (Figure 3F), whereas no such effect was seen in the RPGs, although here we note that the small population of RPGs bearing non-consensus BP sequences leaves the significance of this later result less clear. Nevertheless, within the nRPGs, an additive effect is observed in introns with both non-consensus 5’SS and BP as compared to those with only 1 non-consensus sequence (Figure. 3G).

Together, the previous results suggested to us that RPGs in *S. cerevisiae* may have evolved strong BP, 3’SS, and poly-U tract sequences to facilitate efficient splicing, and we hypothesized that this may be a conserved feature of RPGs due to their critical role in all organisms. To investigate this possibility, we asked about the relative strengths of the 5’SS and 3’SS sequences in three other organisms with well-annotated genomes and well characterized intronomes: the fission yeast *Schizosaccharomyces pombe*, *Drosophila melanogaster* and *Homo sapiens*. For *S. pombe*, we generated PWM scores for the 5’SS and 3’SS as described earlier (see also Methods), whereas for *D. melanogaster* and *H. sapiens* we used previously calculated values. Despite large differences in intron distribution and architecture between *S. cerevisiae* and each of these organisms, a similar trend was seen when comparing RPGs with nRPGs in each of these organisms. Like *S. cerevisiae*, the 3’SS scores for RPGs were significantly stronger in all three organisms as compared with the nRPG scores. (Figure 3H and Table S4). Interestingly we also observed significantly stronger 5’SS scores in *H. sapiens* RPGs compared to nRPGs (Figure 3H).

While these data support the notion that 5’SS and BP sequences are major determinants of 1^st^ step splicing rates, these features alone cannot account for the large differences in 1^st^ and 2^nd^ step rates observed between the RPGs and nRPGs. We hypothesized that there may be cis features of RPGs that distinguish them from nRPGs and contribute to increased 1^st^ and 2^nd^ step splicing rates. Significant differences exist in both intron position and gene structure between RPGs and nRPGs (Figure S4C). Specifically, RPGs have significantly shorter TSS to 5’SS distances and 3’SS to polyadenylation site (PAS) and shorter 3’UTR lengths when compared to nRPGs (Figure S4C). However, RPGs have significantly longer introns compared to nRPGs (Figure S4C), such that there is no significant difference in the unspliced transcript lengths between RPGs and nRPGs (Figure S4C). Within introns, RPGs have significantly longer BP to 3’SS lengths and the nucleotide content in that region is significantly different with RPGs having a higher uridine and lower guanidine and cytidine content (Figure S4C). In contrast, RPGs have a significantly lower fraction of pyrimidines and higher fraction of purines in their full-length intron sequences (Figure S4C).

We then examined these features relative to observed splicing half-lives within the context of the RPG and nRPG classes separately (Figure 4A). Specific to the RPG class, we observed a significant negative correlation between the 3’SS to PAS lengths and the half-lives of the total splicing reaction (Figures 4A and 4B), suggesting that splicing is inefficient when the 3’SS to PAS length is short. Specific to the nRPG class, we found significant negative correlations between the 1^st^ step half-lives and: the length of the 5’UTR, the overall length of the intron, the distance from the 5’UTR to the 5’SS, and the distance between the BP and the 3’SS (Figures 4A and 4C). Together these data suggest that longer distances from the TSS to the end of the intron promote more efficient 1^st^ step splicing in nRPGs. Nucleotide content within the BP to 3’SS region correlated with splicing half-lives in both RPGs and nRPGs. Specifically, the fraction of adenosines in this region showed a significant positive correlation with the 1^st^ step but a significant negative correlation with the 2^nd^ step for both RPG and nRPG classes. The opposite effect was observed for uridine content (Figures 4A and 4D). Surprisingly, within the RPG class, the synthesis rate showed a significant negative correlation with the half-life of the 1^st^ step, but a significant positive correlation with the half-life of the 2^nd^ step (Figure 4A). Neither of these correlations was seen in the nRPG class (Figure 4A).

**Figure 4.**
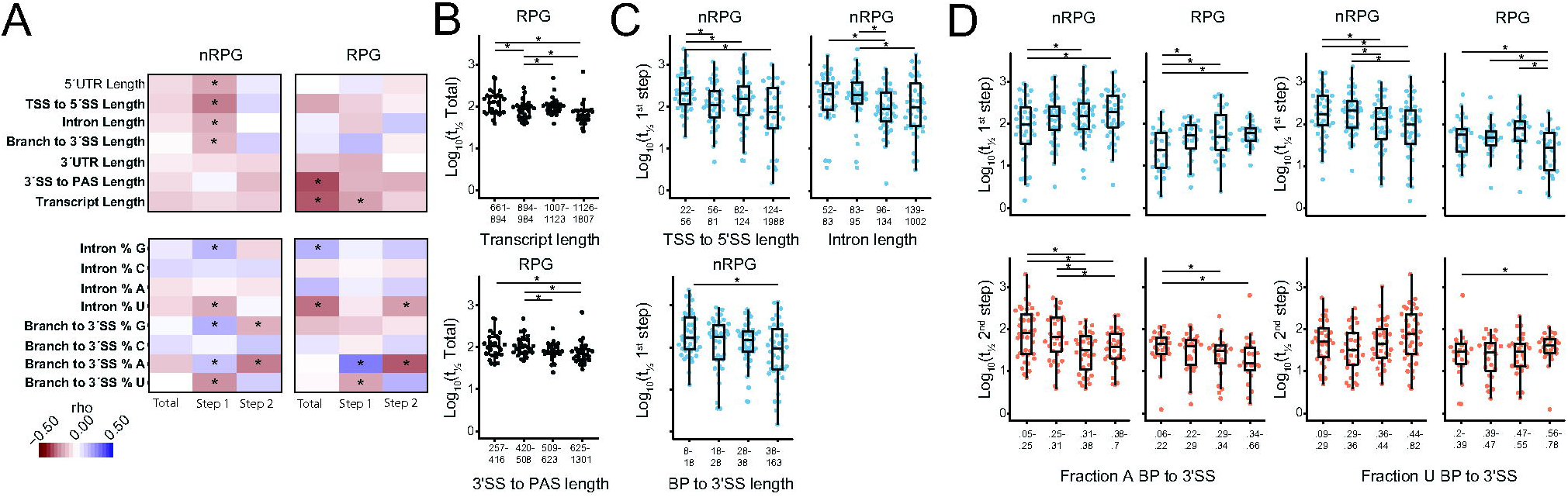
Distinct Transcript Features in RPGs and nRPGs are Correlated with Splicing Rates. (A) Spearman correlations between transcript features and the half-lives for the total, 1^st^ step, and 2^nd^ step of pre-mRNA splicing for RPGs and nRPGs. Bolded features show significant differences between RPGs and nRPGs shown in figure S4 (B) Quartiles of unspliced transcript length and 3′SS to PAS length versus the half-lives for total splicing within RPGs. (C) Quartiles of TSS to 5′SS, intron, and BP to 3′SS lengths versus half-lives of the 1^st^ step of splicing within nRPGs. (D) Quartiles of fraction adenosine in the BP to 3′SS region versus half-lives for the 1^st^ and 2^nd^ steps of splicing for both RPGs and nRPGs. (E) Quartiles of fraction uracil in the BP to 3′SS region versus half-lives for the 1^st^ and 2^nd^ steps of splicing for both RPGs and nRPGs. Statistical significance was calculated with one-sided Mann-Whitney signed-rank test. * p < 0.05.

### A genetic variant reveals expected impacts on the 1^st^ step but unexpected impacts on the 2^nd^

To both confirm the biological significance of our measured rates and assess our ability to detect changes in splicing rates we re-measured global splicing rates in a strain harboring a mutation in the splicing factor Prp2, a DEAH-Box ATPase required for remodeling the spliceosome into a catalytically active state prior to the 1^st^ chemical step. Cells harboring the *prp2-1* allele are inviable at elevated temperatures and show a strong defect in genome-wide pre-mRNA splicing, consistent with its general role in spliceosomal activation. While these cells are viable at room temperature (22°C), their growth is impaired compared to WT, consistent with the strain exhibiting a modest molecular splicing defect even at the viable temperature (Pleiss et al., 2007b). To test the efficiency of 1^st^ step splicing in this strain, we measured splicing rates when it was grown at the permissive temperature of 22°C, the same growth temperature that was used for the WT time course. Consistent with the role of Prp2 in the 1^st^ step of splicing we observed a significantly slower median half-life for the 1^st^ step of splicing in the *prp2-1* strain (286 sec) compared to WT (83 sec) (Figure 5A and Table S5). Surprisingly, we also observed a significantly slower median half-life for the 2^nd^ step of splicing in the *prp2-1* strain (233 sec) compared to WT (41 sec) (Figure 5A). Both trends were observed for the RPG and nRPG classes alike (Figure 5A), however RPGs were significantly more impacted at both steps when compared to nRPGs (Figure 5B).

**Figure 5.**
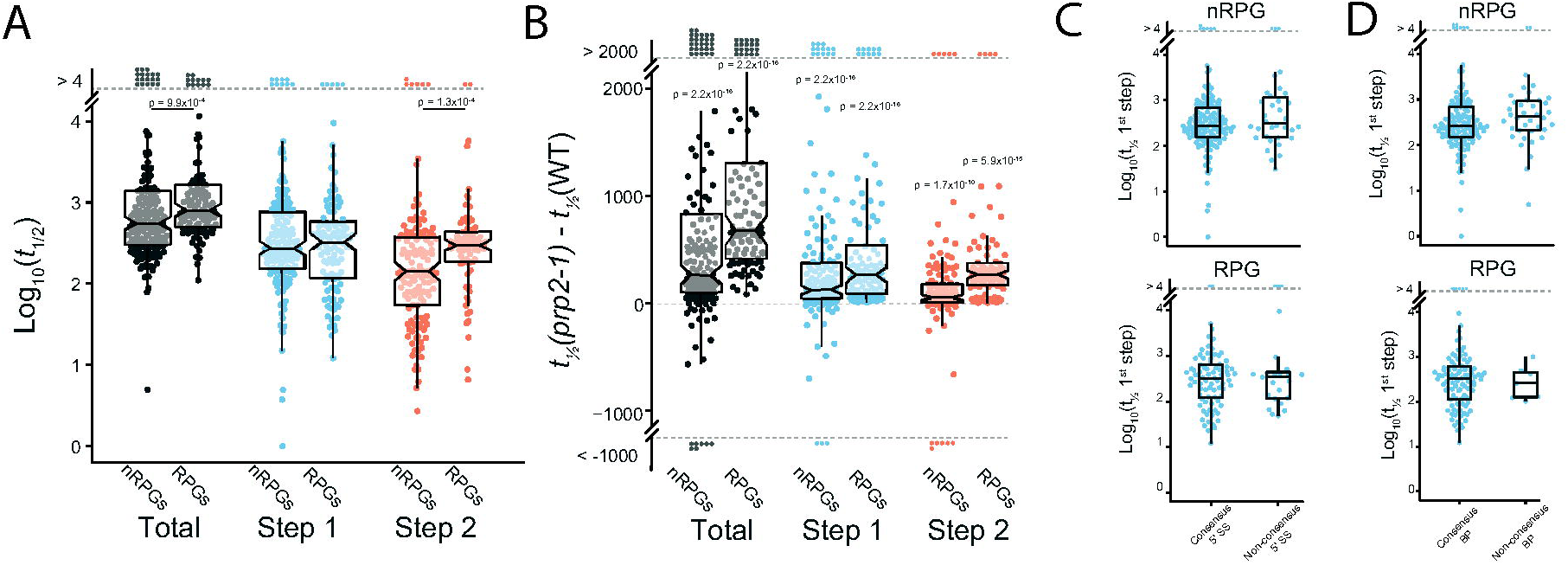
The *prp2-1* allele significantly slows the rate of splicing and differentially impacts RPG and nRPG processing. (A) Distributions of the half-lives for the total (RPG n = 105, nRPG n = 170), 1^st^ step (RPG n = 104, nRPG n = 164), and 2^nd^ step (RPG n = 79, nRPG n = 131) of pre-mRNA splicing in cells harboring the *prp2-1* allele. (B) Distributions of the differences in half-lives between WT and *prp2-1* cells for the total, 1^st^ step, and 2^nd^ step of splicing of RPG and nRPG introns. (C) Comparison of the half-lives for the 1^st^ step of splicing between introns with consensus (RPG n = 80, nRPG n = 130) and non-consensus (RPG n = 24, nRPG n = 32) 5′SS sequences. (D) Comparison of the half-lives for the 1^st^ step of splicing between introns with consensus (RPG n = 86, nRPG n = 126) and non-consensus (RPG n = 7, nRPG n = 38) BP sequences. Statistical significance was calculated with one-sided Mann-Whitney signed-rank test. * p < 0.05

Because Prp2 functions after intron recognition and spliceosome assembly but prior to the 1^st^ catalytic step, we expected that those cis-features which were well correlated with the 1^st^ step rate in WT cells would no longer be determinative of rate when the activity of *prp2-1* became limiting for the 1^st^ step. Consistent with this hypothesis no difference in the half-lives of the 1^st^ step was observed between introns with non-consensus splice site sequences (5’SS or BP) from those with respective consensus sequences in either the RPG or nRPG classes (Figures 5C and 5D). Moreover, previously observed correlations between half-lives and BP, 3’SS, U-tract, and 5’SS scores were no longer apparent in the *prp2-1* strain (Figure 6A). We did, however, observe a significant negative correlation between 1^st^ step half-lives and U-tract scores in nRPGs, albeit much milder than what was observed in WT cells (Figures 3D and 6A). Interestingly, we observed a positive correlation between U-tract scores and total splicing half-lives in RPGs (Figure 6A). In addition, we saw the same effect when we assessed the correlation between U-tract score and the difference in half-lives between WT and *prp2-1* in RPGs, suggesting that RPG introns with stronger U-tract scores are more impacted by the *prp2-1* mutation (Figure 6A).

**Figure 6.**
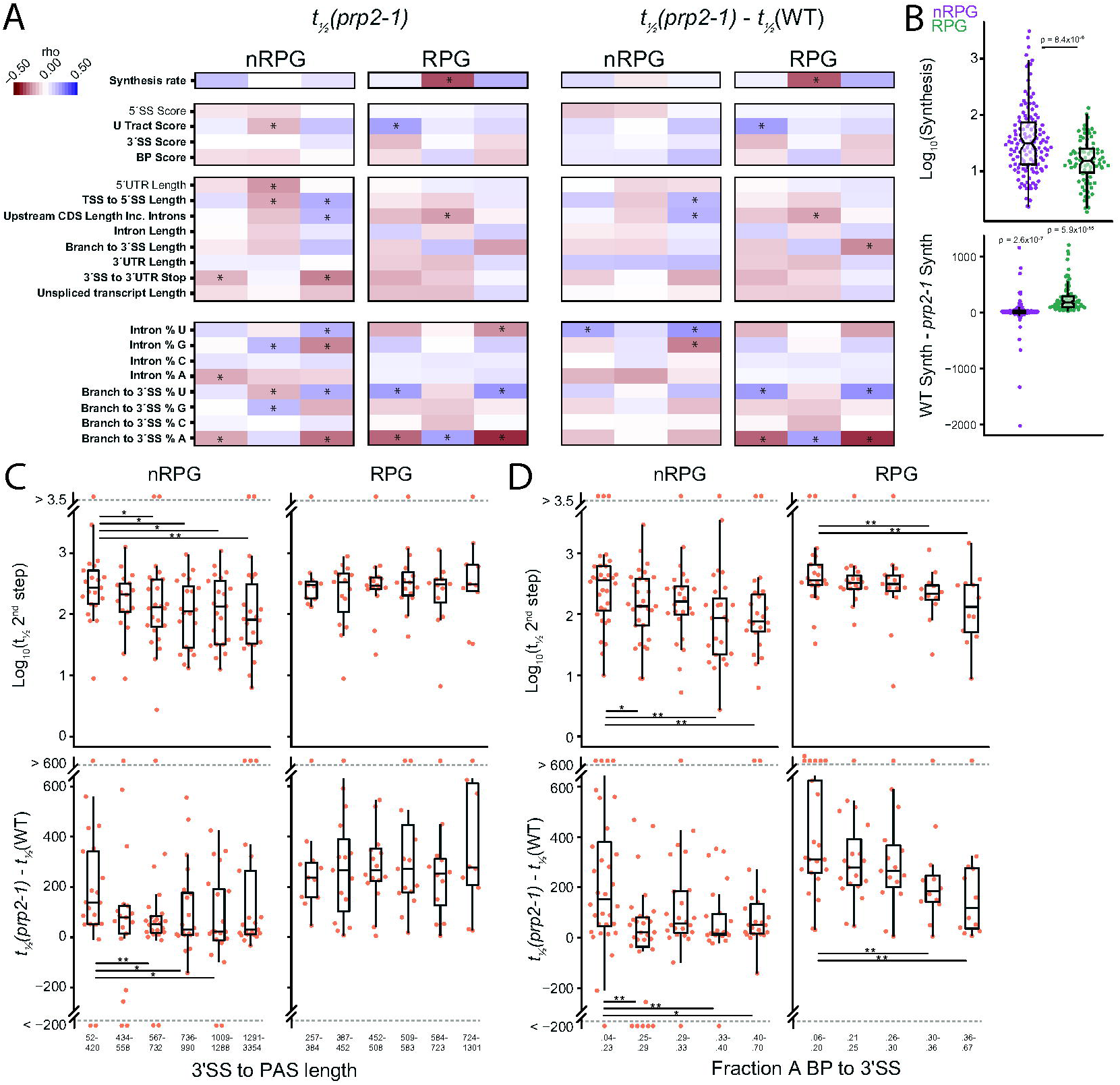
Transcript features correlate with magnitude of *prp2-1* impact on splicing rates. (A) Spearman correlations between transcript features and half-lives and half-live differences between WT and *prp2-1*. Data are partitioned into RPGs and nRPGs. Bolded features show significant differences between RPGs and nRPGs shown in figure S4. (B) nRPG and RPG synthesis rates in *prp2-*1. Synthesis rate difference between WT and *prp2-1.* (C) Sextiles of 3′SS to PAS length in RPGs and nRPGs versus 2^nd^ step half-lives in *prp2-1* and 2^nd^ step half-life differences between WT and *prp2-1.* (D) Quintiles of fraction adenosine in the BP to 3′SS region in RPGs and nRPGs versus 2^nd^ step half-lives in *prp2-1* and 2^nd^ step half-life differences between WT and *prp2-1*. Statistical significance for panel B was calculated with one-sided Wilcoxon signed-rank test. Statistical significance for panels B, C, and D was calculated with one-sided Mann-Whitney signed-rank test. * p < 0.05, ** p < 0.01, *** p < 0.001

Consistent with its slow growth phenotype, the synthesis rates of RPGs in the *prp2-1* strain were significantly slower when compared to their rates in WT cells (Figure 6B). In fact, whereas RPGs showed faster synthesis rates than nRPGs in the WT strain, they showed significantly slower synthesis rates than nRPGs in the *prp2-1*-containing strain. By contrast, nRPG synthesis rates were largely unchanged from WT (Figure 6B). Importantly, a significant negative correlation was observed between the half-lives for the 1^st^ step of splicing and synthesis rates of RPGs. Likewise, the differences in half-lives between WT and *prp2-1* cells for the 1^st^ step of splicing of RPG introns showed a significant negative correlation with synthesis rates. These data indicate that RPGs are significantly repressed via transcription in the *prp2-1* strain, and those transcripts whose synthesis rates are more impacted in this strain are also more impacted at the 1^st^ step of splicing. Interestingly we no longer observed a significant negative correlation between transcript lengths and 1^st^ step rates in RPGs in *prp2-1* (Figure 6A). However, we observed a significant negative correlation between 3’SS to PAS lengths and 2^nd^ step half-lives in nRPGs, suggesting that introns within transcripts with longer 3’SS to PAS lengths undergo the 2^nd^ step faster than those with short 3’SS to PAS lengths (Figures 6A and 6C). No similar correlation was seen in RPGs, however we note that RPGs as a class have significantly shorter 3’SS to PAS lengths compared to nRPGs (Figures 6A and 6C and S4C).

Similar to observations in the WT data, many of the same correlations were observed in the *prp2-1* strain between nucleotide content in the BP to 3’SS regions and 1^st^ and 2^nd^ step half-lives. Specifically, a significant negative correlation was observed between the fraction of adenosines in this region and the 2^nd^ step half-lives in both RPGs and nRPGs (Figures 6A and 6D). Additionally, significant negative correlations were observed between the fraction of adenosines in the BP to 3’SS region and the difference in the 2^nd^ step half-lives between the WT and *prp2-1* strains in both the RPG and RPG classes (Figures 6A and 6D). To a lesser extent, the inverse was observed with the fraction of uridines in this region.

## Discussion

The process of pre-mRNA splicing is pervasive throughout eukaryotes and provides an important control point for regulating gene expression. The capacity of the spliceosome to efficiently process full complement of substrates is poorly understood, yet knowledge of this will be critical for solving problems of both basic and clinical importance. Here we have combined rapid metabolic labeling and first order kinetic modeling with a targeted sequencing approach termed MPE seq to gain high resolution information about the efficiency by which this enzyme processes its full complement of substrates in the budding yeast *S. cerevisiae* (Figures 1A and 1B). While the basic machinery that catalyzes intron removal is highly conserved across eukaryotes, in yeast the intron structures and splicing patterns have greatly simplified over time, facilitating a comprehensive analysis of the splicing targets (Spingola et al., 1999, Wahl et al., 2009). Whereas others have taken a similar approach to understanding this problem in a variety of organisms including yeast, the resolution provided by MPE-seq in the current work enables unmatched resolution of the temporal, global, and chemical aspects of this process (Barrass et al., 2015, Eser et al., 2016, Pai et al., 2017, Wachutka et al., 2019, Windhager et al., 2012, Rabani et al., 2014).

### Splicing efficiency is highly variable across the genome-wide complement of substrates

An important and somewhat unexpected conclusion from the data presented here is that there exists a great variability in the efficiency with which the spliceosome processes its global complement of substrates, even within the relatively simplified system present in budding yeast. In considering this statement, it seems important to acknowledge certain limitations inherent to experiments such as those presented here, in particular the ability to accurately measure very fast and very slow events. Although the time points included in our experiments were selected because they optimized the fraction of splicing events that would be well sampled within our data, there remain some number of events for which the accuracy of our measurements is lower than desired. Nevertheless, the window through which we can robustly measure these rates constitutes a >30-fold difference between the fastest (with half-lives ∼30s in a WT strain) and slowest (with half-lives ∼15m) events (Figure 2A and Table S2). Indeed, our data reveal a continuum of rates for splicing ranging between and beyond these limits, highlighting the dramatic variability of speeds through which different introns transit this process (Figure 2A and Table S2). Importantly, these experiments demonstrate that this variability isn’t the result of variability in just one of the chemical steps, rather significant variation is apparent for both chemical steps of splicing (Figure 2A). While a global view of our data shows that the 2^nd^ step is roughly twice as fast as the first step, suggesting that the 1^st^ step is generally rate limiting, there are many splicing events for which the 2^nd^ step is slower than the 1^st^ step, suggesting splicing has been optimized differently for different substrates (Figures 1C and 2A).

### Cis-elements are important for, but not fully determinative of, splicing efficiency

Importantly, the kinetic measurements reported here are consistent with the long-held idea that splice site sequences play an important role in facilitating splicing efficiency. Indeed, the efficiencies of the 1^st^ chemical step measured here are strongly correlated with the strengths of the splice site sequences, both individually and in composite (Figure 2B-E). Here again, however, it is important to note that while the use of Position Weight Matrix scores enables a powerful approach for comparing the relative activities of these substrates, there are limitations and caveats to such an approach. First and foremost, it is important to emphasize that scores generated in this way are not direct measures of activity *per se*, but rather are measures of frequency; as such a rare but otherwise efficient sequence would appear as low scoring in this approach. Moreover, as noted earlier, the high similarity of sequences found within *S. cerevisiae* introns, particularly at the 5’SS, reduces the capacity of this approach to effectively differentiate between activities. Nevertheless, and these caveats notwithstanding, we note that splice site sequences alone are insufficient to fully the explain complement of 1^st^ step rates determined here. This fact is perhaps best exemplified by the number of introns with fully canonical splice site sequences which nevertheless show disparate splicing efficiencies (Tables S2 and S3). Similarly, the absence of strong correlations between splice site sequences and 2^nd^ step efficiencies highlights the important roles that other substrate features must play in driving splicing rates (Figure 2B).

Beyond the simple global analyses, our analyses of individual variants also provide insights into mechanisms of spliceosomal activation by revealing the differential impact of non-consensus splice site sequences on splicing efficiency (Figures 2B–2E). This was particularly apparent for variants of the 5’SS and their impact on the 1^st^ step of splicing. Among introns with a non-consensus 5’SS, those with a U to A substitution at the 4^th^ position had the slowest 1^st^ step splicing half-lives, whereas those with a C substitution at the 4^th^ position showed no difference from introns with consensus sequences (Figure 2C). The consensus 5’SS in budding yeast is perfectly complimentary to the U1 snRNA, except at the 4^th^ position (Figure 2B). By contrast, the canonical U at this 4^th^ position is complementary to the U6 snRNA in the B^act^ complex (Zhang et al., 2019). Interestingly, while both the A and C mutations result in a mismatch with the U6 snRNA, an A at the 4^th^ position of the 5’SS results in perfect complementarity to the U1 snRNA, whereas a C maintains the mismatch. The observation here that U4A variants are inefficiently spliced is consistent with previous studies that showed that extending the regions of complementarity between the U1 snRNA and the 5’SS beyond the canonical, 6-nucleotide region resulted in inefficient splicing (Staley and Guthrie, 1999). Indeed, the data presented here suggest that increased stabilization of the 5’SS:U1 snRNA duplex even within this 6-nucleotide region may be sufficient to impede the capacity of Prp28 to disrupt this region in the pre-B to B complex transition. It is more difficult to draw conclusions about the importance of the 5’SS:U6 snRNA duplex on the basis of these experiments. Whereas the observation here that U4C variants are spliced with similar efficiency to consensus substrates might mean that the 5’SS:U6 snRNA duplex is tolerant to mismatches, it is also possible that these mis-matches could become rate-limiting under different cellular conditions (Figure 2C). By contrast with the 5’SS sequences, we did not observe a differential impact between the compliment of non-consensus BP sequences on 1^st^ step splicing half-lives (Figure 2D), although low sample numbers could be obfuscating meaningful differences.

The data presented here further reveal the presence of a small, conserved region of uridines just upstream of the 3’SS of some introns which is correlated with faster 1^st^ step splicing (Figures 3D and S3A and S3B). Budding yeast are generally considered to lack a canonical poly-pyrimidine tract (pY) in that no strong pY sequence element is apparent between the BP and 3’SS of most introns, and there is no homolog of U2AF, the protein responsible for pY binding in other eukaryotes (Kupfer et al., 2004, Spingola et al., 1999, Lopez and Séraphin, 1999, Parker and Patterson, 1987). However, *S. cerevisiae* contains a functional equivalent of U2AF^65^, Mud2p, which together with the branchpoint binding protein (BBP) binds to the BP and downstream region in the initial steps of spliceosome assembly (Abovich et al., 1994). Our data suggest that this U-tract region may function similarly to the pY tract in other eukaryotes.

Beyond the U-tract element, our data also suggest that overall nucleotide content within the BP to 3’SS region may be important for splicing efficiency at both steps. Introns with high A content in this region tend to be processed slowly through the 1^st^ step but fast through the 2^nd^ step, whereas the inverse was observed with U content (Figures 4A and 4D). To be sure, it is difficult to deconvolute the presence of a strong U-tract element from the more general property of A and U content within the region, leaving it unclear the mechanisms by which nucleotide content within this region might influence splicing rates. Nevertheless, it is important to note that throughout the splicing cycle this region is in close proximity to many spliceosomal proteins which might impart selective behavior on select transcripts based on these properties (Will and Lührmann, 2011, Fica and Nagai, 2017, Wahl et al., 2009). Similarly, it is well established that secondary structures can impact splicing efficiency, and these correlations may reflect increased or decreased capacity for formation of such structures (Warf and Berglund, 2010, Barrass et al., 2015).

Importantly, our work with a strain containing a conditional genetic variant of the essential splicing factor Prp2 further reinforces the role of cis-regulatory elements in splicing activity. Prp2 acts just prior to the 1^st^ chemical step of splicing after recognition of the 5’SS and BP sequences and assembly of the spliceosome (Wahl et al., 2009). As expected, we observed a global slowing of 1^st^ step splicing rates (Figures 5A and 5B). However, consistent with the late action of Prp2 relative to splice site recognition, we no longer observed a correlation between splice site strength and 1^st^ step rates, suggesting that the rate limiting step is Prp2 function rather than splice site recognition (Figures 5C and 5D).

### Evolution has tuned intronic and genic features to facilitate splicing of classes of transcripts

The relative contribution of splice site sequences in splicing efficiency is perhaps most notable when considering the processing of different classes of transcripts. It has long been known that the ribosomal protein genes (RPGs) constitute a unique category within the subset of intron-containing genes in budding yeast. Whereas only ∼5% of all genes contain introns, nearly 70% of RPGs contain them. Indeed, because of the high transcriptional frequency of RPGs it has been noted that nearly 35% of all transcriptional flux requires processing by the spliceosome, in spite of this low population of introns within the genome (Warner, 1999). Moreover, it was long ago noticed that certain properties of RPG introns were distinct from those in nRPGs, notably that intron lengths are markedly longer in the RPGs (Spingola et al., 1999, Parker and Patterson, 1987). Here we further note that the combination of the long intron lengths but relatively short coding and UTR lengths of RPG results in transcripts whose overall unspliced lengths are virtually indistinguishable from nRPGs, but importantly where the 3’SSs are both farther from the transcription start sites (TSSs) and closer to the cleavage and polyadenylation sites (PAS) (Figure S4). Moreover, we also show here that RPGs as a class are characterized by stronger scoring splice site sequences than nRPGs at the 5’SS, BP, U-tract, and 3’SS regions in budding yeast introns (Figure S4). Importantly, while far from a rigorous evolutionary analysis, an examination of three divergent organisms for which robust genome-wide information about intronic sequence elements is available suggests that strong splice site sequences may be an evolutionarily conserved property of RPGs.

Our data further show that RPG introns transit through both chemical steps of splicing faster than nRPG introns (Figure 3A). Surprisingly however, and somewhat paradoxically, we did not detect any significant correlation between the splicing rates for RPGs and any of the cis-features within their introns. By contrast, within the relatively weaker scoring nRPG introns, 1^st^ step rates were strongly correlated with both 5’SS and BP scores (Figures 3D–3E). Additionally, the combination of nRPG introns with non-consensus 5’SS and BP sequences showed an additive impact on 1^st^ step half-lives (Figure 3G). Moreover, U-tract score correlated strongly with 1st step half-lives in nRPGs, whereas no significant correlation was observed in RPGs.

How then to understand the presence of such strong splice site sequences within the RPG introns, and the high efficiency with which they are processed, but the apparent insensitivity of the spliceosome to the non-consensus variants within this class? As described below, we suggest that the data presented here are most consistent with a model wherein the overall architecture of the ribosomal protein genes has been tuned to enable efficient splicing, in particular by connecting their splicing to transcription, such that under optimized growth conditions the spliceosome is insensitive to suboptimal splice site sequences. Presumably the strong splice site features remain positive determinants of efficient RPG splicing under suboptimal growth.

Two main observations drive this hypothesis. First, the data presented here show that transcript length upstream of the 3’SS is a general effector of 1st step splicing rates, suggesting that RPGs may have selected for long introns to maximize 1st step splicing efficiency (Figures 4A and 4C). Mechanistically, increased length may provide additional time for association of the U1 snRNP and other early spliceosome components with the nascent transcript and transcription machinery, allowing for quicker progression through the splicing cycle once the full intron has been synthesized. Second, RPGs as a class are subject to higher transcription frequency, measured here as synthesis rates (Figure 3B). While, it remains poorly understood how transcriptional activity and splicing rates are coupled, over the past decade increasing evidence has accumulated pointing to a role for liquid phase separation in gene expression, in particular during the early stages of transcription (Boehning et al., 2018, Lu et al., 2019, Harlen and Churchman, 2017, Hnisz et al., 2017). We propose that the high transcriptional activity of RPGs in WT cells is accompanied by liquid phase-separated compartments containing higher concentrations of splicing factors leading to fast splicing rates relative to nRPGs. A comparison of splicing rates in WT cells versus those harboring the *prp2-1* variant further supports this idea. Here, a strong correlation is observed between the change in transcriptional frequency and the change in splicing rate: as transcription rate slows, so does splicing. Decreased transcription is presumably accompanied by a reduction of these phase-separated compartments, leading to slower splicing.

### Splicing is fast, but occurs over the length of the transcript, completing when polymerase is thousands of bases downstream

While the experiments presented here evaluate the kinetic properties of splicing intermediates, and as such do not directly evaluate the position of RNA polymerase with respect to this process, as noted in the previous section much can be inferred from these data about the relationship between splicing and transcription. Importantly, while the current inconsistencies within the literature regarding the relationship between polymerase location and splicing status might have been discounted as merely species-specific differences in the pathways of spliceosome assembly and chemistry, the data presented here for budding yeast are more consistent with the observations from higher eukaryotes wherein spliceosome assembly is presumed to occur in a co-transcriptional manner across the length of the downstream exon, with the chemistry of splicing completing at a position thousands of nucleotides downstream of the intron. Three different aspects of our data drive this conclusion.

First and foremost, the splicing half-lives measured here are on the order of minutes, with a median half-life of just over two minutes. While estimates of the elongation rate of RNA polymerase vary by organism and growth condition, it is generally accepted that this occurs at a rate of roughly 1-4 kilobases per minute (Mason and Struhl, 2005, Oesterreich et al., 2011). To be sure, elongation rates are likely to be on the slower end of these estimates for yeast growing at 22°C, but even at the low end of this spectrum the polymerase is expected to be more than two kilobases downstream of the intron when the median splicing event completes. Moreover, while RNA polymerase does not transcribe at a uniform rate through a gene and transient pausing has been observed, the lifetime of transcriptional elongation pauses are measured in seconds, or fractions thereof, and as such are unlikely to significantly impact the location of the polymerase relative the times measured here (Oesterreich et al., 2011, Kwak et al., 2013).

Second, as noted above, the overall splicing rates presented here show a strong correlation with splice site strengths, cis-elements long established to influence splicing efficiency (Figures 2B-E). The poor annotations of global splice site sequences, particularly BPs, in most higher eukaryotes has presumably precluded a careful assessment of the correlations between these features and polymerase location in earlier studies. However, we note that we detect no correlation between splice site strengths and the median splice distance in the yeast studies that report the completion of splicing in close proximity to the 3’SS (Oesterreich et al., 2016).

And finally, the combination of our measurements in both WT and *prp2-1* cells reveals an important correlation between the length of the transcript between the 3’SS and the PAS and the efficiency of splicing, wherein the longer the distance the more efficient the splicing. Such an observation would not be expected if splicing was completed very rapidly in relation to transcription of the 3’SS. Rather, the simplest explanation for this observation is that the splicing process is facilitated by its connection with the polymerase, and the longer the region downstream of the intron but upstream of the cleavage site the longer-lived is the connection with the polymerase. Indeed, our observation that RPG transcripts with long 3’SS to PAS lengths are spliced significantly faster than short ones suggests that the more time in which the nascent RNA is engaged in transcription the more efficient the splicing becomes (Figures 4A and 4B). Moreover, while no correlation was detected between 3’SS to PAS length and splicing of nRPG introns in WT cells, the observation in the *prp2-1* strain that RPG introns as a whole and the subset of nRPG introns with short 3’SS to PAS lengths were significantly slowed at the 2nd step compared to WT again points to this important relationship (Figure 6C). Specifically, nRPG introns with 3’SS to PAS lengths of less than roughly 750nt were slowed in a length dependent manner, while those with 3’SS to PAS lengths greater than 750nt were not (Figure 6C). Importantly, nearly 90% of RPGs have 3’SS to PAS lengths less than 750nt.

## STAR Methods

### RESOURCE AVAILABILITY

#### Lead Contact

Further information and requests for resources and reagents should be directed to and will be fulfilled by the Lead Contact, Jeffrey A. Pleiss

#### Materials Availability

This study did not generate any unique reagents

#### Data and Code Availability

Raw sequencing data is available through…. Python and r code used for processing and analyzing data in this study can be found at https://github.com/mgildea87/4tu_MPEseq

### EXPERIMENTAL MODEL AND SUBJECT DETAILS

#### Strain growth and 4tu time course

BY4741 (WT) and *prp2-1* cells were streaked onto YPD plates from glycerol stocks stored at -80°C and grown for roughly 48 hours at 30°C. Five colonies were inoculated into 50 mL of liquid YPD medium in 250mL flasks and grown in a shaking incubator at 30°C overnight. Triplicate cultures were started for each strain by back diluting the saturated overnight culture to an OD_600_ of 0.1 in 600mL of fresh complete synthetic medium containing 178 μM uracil in a 2.8L flask. Cultures were grown in a shaking incubator at 30°C. After one doubling, cultures were shifted to 22.5°C for an additional two doublings. 4-thiouracil was added to a final concentration of 500 μM to the log phase cells to start the time course. At each time point, 50mL of culture was extracted, vacuum filtered, and flash frozen in liquid nitrogen. Extracted cells were stored at -80°C. Time 0 samples were collected before addition of 4-thiouracil.

### METHOD DETAILS

#### In vitro transcription of 4sU labeled spike-ins

PCR primer sets were designed to amplify five *S. pombe* genes (Table S6). The T7 promoter sequence was appended with PCR such that the resulting amplified DNA could be directly used in *in vitro* transcription. After an initial denaturation for 30 seconds at 98°C, 40 cycles of PCR were performed on a 200 μL reaction for each primer set, including 400 ng of *S. pombe* genomic DNA as template, Phusion Each cycle consisted of 10 seconds at 98°C, 20 seconds at 62-65°C, followed by 30 seconds at 72°C. This was followed by a final extension for 5 minutes at 72°C. All reactions were run on 0.8% agarose gels and bands were gel purified (Invitrogen Purelink) followed by ethanol precipitation. *In vitro* transcription was performed using the NEB HiScribe T7 high yield synthesis kit following the manufacturers protocol using 400 ng of each DNA template in 20 μL reactions. RNA was 4-thiouracil labeled by including 4-thiouridine-5’-triphosphate (4tUTP) at 1:3 with uridine-5’-triphosphate (UTP). 1:3 was empirically chosen because it afforded maximum purification efficiency of 4sU labeled spike-in transcripts compared to other tested ratios (data not shown). RNA was phenol chloroform purified following the manufacturers protocol. The length of 4sU labeled spike-in transcripts was assayed using the 2100 Bionalyzer (Agilent). The vast majority of transcripts were the expected full length (data not shown). 4sU labeled spike-ins were pooled at equal mass and stored at -80°C in 10 mM Tris-HCl pH 7.4, 1 mM EDTA.

#### RNA Extraction

RNA was extracted by adding 2 mL acid phenol (pH 5.3) to each cell pellet followed by 2 mL of AES buffer (50 mM sodium acetate pH 5.3, 10 mM EDTA, 1% SDS). Samples were incubated for seven minutes at 65°C with periodic mixing via vortexing. This was followed by a five minute incubation on ice. Samples were transferred to 15 mL PLG tubes and spun for five minutes at 3000g. 2 mL of phenol:chloroform:IAA (25:24:1) was added to the aqueous phase in each tube and samples were mixed and centrifuged for five minutes at 3000g. To remove excess phenol, one more extraction performed with 2 mL chloroform was performed. RNA was then isopropanol precipitated by addition of 200 μL 3 M Sodium acetate (pH 5.3) followed by 2.5 mL isopropanol. RNA was pelleted, washed twice with 70% ethanol, dried, and dissolved in 10 mM Tris-HCl (pH 7.4), 1 mM EDTA.

#### Biotin coupling to 4-thiouracil labeled RNA

For each sample, 400 μg of total RNA was mixed with 2.5 ng of 4-thiouracil labeled spike-in mix. Biotin coupling was performed following a previously published protocol(Dolken et al., 2008). 100 μL of 100 mM Tris-HCl (pH 7.4), 10 mM EDTA was added to each sample followed by DEPC water up to 800 μL. This was followed by the addition of 200 μL of HPDP-biotin (1 mg/mL in dimethyl formamide). Each sample was mixed well and placed on a rotator at room temperature in the dark for three hours. Two 1 mL chloroform extractions in 2 mL PLG tubes were performed to remove excess unreacted HPDP-biotin. Biotin coupled RNA was isopropanol precipitated and RNA pellets were dissolved in 100 mL of 10mM Tris-HCl (pH 7.4), 1 mM EDTA.

#### Biotin purification

Biotin purification was performed according to a previously published protocol(Dolken et al., 2008). 100 μL of streptavidin beads (Dynabeads C1 Invitrogen) per sample were transferred to tubes and placed on a magnetic stand for one minute. The supernatant was aspirated, and the tubes were removed from the magnetic stand. The beads were washed by resuspending the pellet in 1 mL of bead wash buffer (100mM Tris-HCl pH 7.4, 10mM EDTA, 1M NaCl, 0.1% Tween) followed by mixing via pipetting. The samples were placed back on the magnetic stand for 1 min and the buffer was aspirated. Washing was repeated 2 more times for a total of 3 washes. Beads were resuspended in a 100 μL per sample volume of bead binding buffer (10 mM Tris-HCl pH 7.4, 1 mM EDTA, 2 M NaCl, 0.1% Tween) and mixed well by pipetting. 100 μL of resuspended beads were added to each biotin coupled RNA sample and placed on a tube rotator in the dark for 30-min at room temperature. Beads were washed 3 times with bead wash buffer pre-warmed to 65°C and three additional washes with room temperature bead wash buffer. To elute RNA, beads were resuspended in 100 μL of freshly prepared 100 mM DTT. Samples were mixed well and incubated at room temperature for 2-min. Eluted RNA was aspirated and transferred to a new tube. The elution was repeated for a total of two elutions. Each RNA sample was purified by adding 1400 μL of binding buffer (2 M Guanidinium-HCl, 75% isopropanol), mixed well by vortexing, and transferred to Zymo-spin I columns. Samples were spun at 14k x g for one minute. This was followed by 2 washes with column was buffer (10 mM Tris-HCl pH 8.0, 80% ethanol). Columns were dried by spinning at 14K x g for an additional one minute. Purified RNA was eluted by adding 16 μL DEPC water. RNA was collected by spinning at 14k for one minute. RNA concentration was assayed using Qubit.

#### MPE-seq library preparation and sequencing

MPE-seq libraries were prepared as previously described with a few key differences(Gildea et al., 2019). 14 μL of each 4-tu purified RNA sample was included in the reverse transcription (RT) reaction. Five additional MPE-seq RT primers targeting the five spike-in RNAs were added to the *S. cerevisiae* pool at equimolar concentration (80 nM) (Table S6). Libraries were barcoded via PCR amplification and pooled. Paired-end sequencing was performed on the NextSeq 500 (Illumina) with a read 1 (P5) length of 61-bp and a read 2 (P7) length of 17-bp. Sequencing was performed by the BRC Genomics Facility at Cornell University.

#### MPE-seq data analysis

##### Read processing and alignment

Reads were demultiplexed by the sequencing center. First, overall read quality was assessed by running FastQC on each fastq file(Andrews, (n.d)). Illumina sequencing adapter sequences were trimmed from reads using fastp (version 0.20.0) with the following parameters --disable_trim_poly_g -- adapter_sequence CTGTCTCTTATACACATCT --adapter_sequence_r2 CTGTCTCTTATACACATCT(Chen et al., 2018). Reads were aligned to the yeast genome (reference genome assembly R64-1-124(Engel et al., 2014)) with STAR (version 2.7.2b) with the following parameters --clip5pNbases 7 0 --peOverlapNbasesMin 5 --peOverlapMMp 0.1 -- outFilterMultimapNmax 1 --alignIntronMin 10 --alignIntronMax 1100 --outSAMattributes All -- runThreadN 8 --outSAMunmapped Within KeepPairs --alignSJoverhangMin 3 -- alignSplicedMateMapLmin 3 --alignMatesGapMax 3000 --alignEndsType EndToEnd(Dobin et al., 2013). Only concordantly aligned reads were considered for further analysis accept for calculation of the spike-in normalization factor wherein total on target reads were used (concordant + non-concordant).

#### Estimating splice isoform abundance

Reads derived from targeted spliced transcripts were identified and counted based on the presence of an ‘N’ in their CIGAR string and an extension of at least 3 bases across the splice junction into the upstream exon. Reads derived from unspliced transcripts were identified by a concordantly mapped read pair in which read 1 aligned to a primer target site and extended into an intron and was on the appropriate strand. Each intron was considered individually. For example, if a read pair aligned to 2 or more introns or splice junctions within a multi-intronic gene, only the targeted intron or splice junction was counted based on the position of read 1. Unspliced reads were further partitioned into lariat intermediate, pre-1^st^ step, and ambiguous unspliced based on the position of read 2. Lariat intermediate reads were those whose 1^st^ mapped base of read 2 aligned within an 8 base pair window around the annotated branch point adenosine. This window used was 5 bases downstream and 3 bases upstream of the annotated branchpoint adenosine. Ambiguous reads were those whose 1 ^st^ mapped base of read 2 aligned downstream of the lariat intermediate window and the 3’SS. Pre -1^st^ step reads were than quantified by subtracting the sum of lariat intermediate and ambiguous unspliced from total unspliced reads. Finally, ambiguous reads were assigned to lariat intermediate or branched based on the ratio of unambiguous lariat intermediate to pre-1^st^ step reads. To remove reads derived from primers not extended by RT, read pairs with insert sizes less than 33nt were discarded. Isoform quantification was performed using a custom python script that can be found in the Data and Code Availability section.

#### Read normalization

To monitor changes in absolute abundance of each RNA isoform over time read counts were normalized to *S. pombe* spike-in counts.

1. For each spike-in *k* in each replicate time course *j*, for each time point *t*. The fraction spike-in counts *A* of total counts:

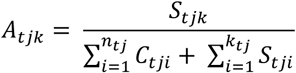 Where *k* = 1-5 for each spike-in, *j* = 1-3 for each time course replicate, *S* = spike-in count, *C* = target intron counts, and *n* = intron targets.
2. We then computed the ratio *B* of each *A* to the *A* of time point 0 *t*_0_ in each replicate time course for each spike-in:

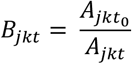
3. To remove outliers (spike-in counts that behaved far from the norm) we combined *B* values across replicates *j* and spike-ins *k* for each time point *t* (15 values/time point). Outliers were identified as values outside of the 1.5 x IQR (identified with the boxplot.stats() function in r) and removed. In the WT data 10 of 150 poorly behaved values were removed. 9 values were removed from the *prp2-1* data.
4. The scaling factor *F* was computed for each time point *t* by computing the geometric mean of *B* for the 5 spike-ins and 3 replicates:

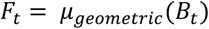
5. To normalize read counts *C* for each intron target *n* in each replicate *j* in each time point *t*, read counts were multiplied by the scaling factor *F*:

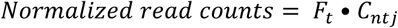

##### Background correction

To estimate and correct for purification of unlabeled RNA during the 4 -tu purification, samples were taken before the addition of 4-tu to the culture media. The mean normalized counts for total unspliced, lariat intermediate, and pre 1^st^ step isoforms for each target were calculated across the time 0 samples and were subtracted from the corresponding counts in each other time point.

##### Model Fitting

The abundance of labeled RNA increases with time primarily due to transcription however, the cells division rate also increases labeled RNA abundance. However, the replication rate of WT and *prp2-1* cells was roughly 100 times slower than the estimated splicing rates. This would be a constant correction across all splicing rates and would only minimally change the rate estimates. For this reason, we did not adjust for growth rate in our model.

##### Total splicing rate model

To model the total rate of splicing, a first order model was fit to background corrected normalized total unspliced counts to estimate the synthesis rate *k_synth_* and total splicing rate *k_splicing_*:

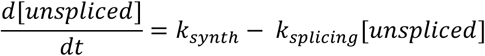

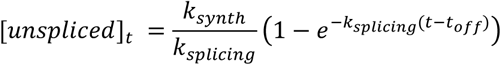

The model was fit to the composite data across the three replicates using non-linear least squares with weights equal to 1/t^2^. This was done to place more weight on earlier time points that generally contain a large portion of the approach to equilibrium curve. The models were fit in r using the nlsLM() function in the minpack.lm package. Several samples behaved far from expected consistently across all targets and were removed from the total splicing rate and further models. 4 of 30 samples were removed from WT and 3 of 30 samples were removed from *prp2-1*.

##### Time offset

When 4-tu is added to a culture of cells, it must be transported into the cell and processed before it becomes available to the transcription machinery in the nucleus. This lag time (*t_off_*) was estimated by fitting the above described model to pre-1^st^ step counts and allowing it to estimate *t_off_* in addition to the other parameters for every intron. To estimate total splicing rates, the modeling procedure was repeated with total unspliced counts using the median *t_off_* as a fixed parameter. *t_off_* was 105s and 182s for WT and *prp2-1*, respectively.

##### Individual step coupled splicing rate model

The abundance of pre 1^st^ step counts [*pre* 1*st step*] over time is influenced by *k_synth_* and decay via the *k*_1*st*_. The abundance of pre 1^st^ step counts over time was modeled using a simple 1^st^ order model as follows:

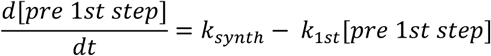

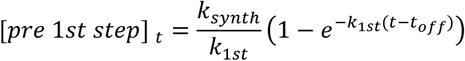

The abundance of lariat counts over time are influenced by *k_synth_*, *k*_1*st*_ and *k*_2*nd*_ The abundance of lariat intermediate counts over time was modeled using a 2 consecutive 1^st^ order reactions model described below:

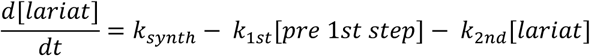

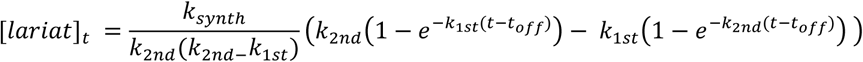

We assume that the same *k_synth_*, *k*_1*st*_ influence pre 1^st^ step and lariat intermediate counts for a given intron. As such, we estimated *k_synth_*, *k*_1*st*_ and *k*_2*nd*_ by simultaneously fitting both pre 1^st^ step and lariat intermediate models with *k_synth_* and *k*_1*st*_ as shared parameters. The models were fit to the composite data across all 3 replicates. As before, NLS was used with weights as described above. 90% confidence intervals for parameter estimates were calculated using the confint2() function in r and reported in the supplemental table (Tables S2, 5) As described, *k_synth_* was estimated in both the total splicing rate and individual step coupled models. Both synthesis rate estimates correlated very well and *k_synth_* from the total splicing rate model was used in synthesis rate analysis throughout this paper.

#### Splice site scores

For *S. cerevisiae* splice site scores (5’SS, BP, 3’SS region, 3’SS, and U-tract) position weight matrices were generated. Mononucleotide probabilities across all introns were used to generate scores and background nucleotide probabilities were calculated from the combination of all intron sequences (Table S3). Composite splice site scores were calculated by adding 5’SS, BP, and 3’SS region scores for each intron (Table S3). *S. pombe* 5’SS and 3’SS scores were calculated using the Burge lab’s MaxEntScan tool’s maximum entropy model(Yeo and Burge, 2004). *H. sapiens* and *D. melanogaster* 5’SS and 3’SS scores were obtained from Drexler H., *et al*(Drexler et al., 2020). Sequence logos were generated from mononucleotide position probability matrices using the ggseqlogo package in R (Figures 2 and S3A and S4B)(Wagih, 2017)

##### Intron and transcript annotations

*S. cerevisiae* Intron, open reading frame, and function (RPG or nRPG) annotations were extracted from the .gff feature file associated with the current genome release (Engel et al., 2014)(R64-2-1 downloaded from *Saccharomyces* Genome Database)(Engel et al., 2014). TSS and PAS annotations were extrapolated from published TIF-Seq median 5’UTR and 3’UTR lengths(Pelechano et al., 2013). *S. pombe* intron annotations and splice site sequences were extracted from the .gff3 file supplied by pombase and the current genome release(Lock et al., 2018, Wood et al., 2002)

### QUANTIFICATION AND STATISTICAL ANALYSIS

Statistical analysis, quantification approaches, and RNA processing rate modeling associated with sequencing data are presented in the Method Details section. Sequencing data processing and transcript/splice isoform quantification were performed using the software described above along with custom python scripts that can be found in the Data and Code Availability section. RNA processing rate modeling was implemented in r. Code for modeling procedure can be found in the Data and Code Availability section. All statistical analyses mentioned in figure legends throughout the manuscript were performed in r.

### KEY RESOURCES TABLE

| REAGENT or RESOURCE | SOURCE | IDENTIFIER |
| --- | --- | --- |
| Chemicals, Peptides, and Recombinant Proteins |  |  |
| Complete synthetic media (powder) | Sunrise Science Products | cat# 1001-100 |
| 4-thiouracil (4tu) | Santa Cruz Biotechnology | Cat# sc-256832, cas<br>591-28-6 |
| Phusion polymerase | NEB | M0530S |
| 4-thiouridine-5'-triphosphate (4tUTP) | TriLink Biotechnologies | Cat# N-1025 |
| Acid-phenol:chloroform, pH 4.5 (with IAA, 125:24:1) | Invitrogen | Cat# AM9722 |
| Phenol:chloroform IAA pH 7.9 (25:24:1) | Invitrogen | Cat# AM9732 |
| Chloroform | Fisher Scientific | Cat#<br>AC383760010,<br>cas# 67-66-3 |
| MaXtract high density tubes 15mL (PLG) | Qiagen | Cat# 129065 |
| MaXtract high density tubes 2mL (PLG) | Qiagen | Cat# 129056 |
| EZ-Link HPDP-Biotin | ThermoFisher Scientific | Cat# 21341 |
| Dynabeads MyOne streptavidin C1 | ThermoFisher Scientific | 65002 |
| Dimethyl formamide (DMF) |  |  |
| Dithiothreitol (DTT) |  |  |
| Guanidinium-HCl |  |  |
| Zymo-spin I columns | Zymo Research | C1003-250 |
| Critical Commercial Assays |  |  |
| PureLink quick gel extraction kit | ThermoFisher Scientific | Cat# K210012 |
| HiScribe T7 high yield RNA synthesis kit | NEB | E2050S |
| Bioanalyzer RNA 6000 nano kit | Agilent | 5067-1511 |
| Qbit dsDNA HS assay kit | ThermoFisher Scientific | Q32851 |
| Deposited Data |  |  |
| Experimental Models: Organisms/Strains |  |  |
| *S.cerevisiae*: BY4741 (WT) | ? | ? |
| *S.cerevisiae*: *prp2-1* | Vijayraghavan, U., et al.(Vijayraghavan and Abelson, 1989) | ? |
| Oligonucleotides |  |  |
| All oligo sequences listed (**Table S6**) | This study | N/A |
| Software and Algorithms |  |  |
| FastQC version 0.11.5 | Andrews, S., et al.(Andrews, (n.d)) | https://www.bioinformatics.babraham.ac.uk/projects/fastqc/ |
| Fastp version 0.20.0 | Chen, S., et al.(Chen et al., 2018) | https://github.com/OpenGene/fastp |
| STAR version 2.7.2b | Dobin, A., et al.(Dobin et al., 2013) | https://github.com/alexdobin/STAR |
| Samtools version 1.6 | Li H., et al.(Li et al., 2009) |  |
| Pysam version 0.15.1 |  | https://github.com/pysam-developers/pysam |
| HTSeq version 0.6.1 | Anders S., et al.(Anders et al., 2015) |  |
| R studio version 1.0.153 |  | Rstudio.com |
| R version 4.0.2 |  |  |
| Python version 2.7.17 |  |  |
| MaxEntScan | Yeo G., et al(Yeo and Burge, 2004) | http://hollywood.mit.edu/burgelab/maxent/Xmaxentscan_scoreseq.html |
| ggseqlogo version 0.1 | Wagih, O., et al.(Wagih, 2017) | https://github.com/omarwagih/ggseqlogo |
| ggplot2 version 3.3.2 | Wickham H.(Wickham, 2016) | https://ggplot2.tidyverse.org |

## SUPPLIMENTAL INFORMATION

Table S1. Key statistics of 4su labeled RNA purification and MPE-seq libraries sequenced in this study, related to STAR methods.

Table S2. WT splicing half-life estimates related to Figures 2–6. Splicing half-life estimates from the total splicing rate and individual step coupled splicing rate models. Upper and lower 90% confidence intervals are included.

Table S3. *S. cerevisiae* cis transcript features related to Figures 2–6.

Table S4. *S. pombe* 5’SS and 3’SS scores related to Figure 3.

Table S5. *prp2-1* splicing half-life estimates related to Figures 5–6. Splicing half-life estimates from the total splicing rate and individual step coupled splicing rate models. Upper and lower 90% confidence intervals are included.

Table S6 Oligonucleotide sequences used in this study, related to STAR methods. Includes oligonucleotide primer sequences for generation of IVT spike-ins and all RT primers used to prepare MPE-seq libraries.

**Figure S1. 4su labeling purifies nascent RNA. Related to Figure 1.**

(A) Mass of 4su labeled RNA purified versus time after addition of 4tu to the media. RNA mass is represented as the mean across 3 replicates for each time point. Error bars are 1 standard deviation.

(B) Mean splice index (SI) versus time after addition of 4tu to the media. SI is defined as total unspliced reads divided by total spliced reads for each intron. SI is represented as the mean across 3 replicates for each time point.

**Figure S2. 1^st^ and 2^nd^ step rates are not correlated. Related to Figure 2.**

(A)Log_10_ transformed 1^st^ step half-lives versus Log_10_ transformed 2^nd^ step half-lives. Spearman correlation coefficient is included (ρ)

**Figure S3.U-tract strength correlates with 1^st^ but not 2^nd^ step rates. Related to Figure 3.**

(A) Sequence logo generated from all analyzed introns for a region around the 3’SS. U-tract and 3’SS portions used for splice site score calculations are highlighted.

(B) Comparison of U-tract and 3’SS scores with 1^st^, and 2^nd^ step rates (see also Table S3). Spearman correlation coefficients (ρ) and associated p-values (p) are included. Contour lines were drawn from 2d kernel density estimation implemented by the stat_density_2d() function in the R package. Color fill gradient corresponds to density level. Red text and highlighted axes indicate significant correlations (p < 0.05).

**Figure S4. 1^st^ and 2^nd^ step rates are not correlated. Related to Figure 2.**

(A) Log_10_ transformed 1^st^ step half-lives versus Log_10_ transformed 2^nd^ step half-lives for RPGs and nRPGs. Spearman correlation coefficients (ρ) and associated p-values (p) are included. Red text and highlighted axes indicate significant correlations (p < 0.05).

(B) Sequence logos generated from RPG or nRPG introns for a region around the 3’SS. U-tract and 3’SS portions used for splice site score calculations are highlighted.

(C) Boxplots comparing transcript features between nRPG and RPGs. Statistical significance was calculated with one-sided Mann-Whitney signed-rank test and associated p-values (p) were included were p < 0.05.

## Notes

### Competing Interest Statement

The authors have declared no competing interest.

