## Supplementary figures and images for "Transcript-specific determinants of pre-mRNA splicing revealed through *in vivo* kinetic analyses of the 1^st^ and 2^nd^ chemical steps"

### Figure S1

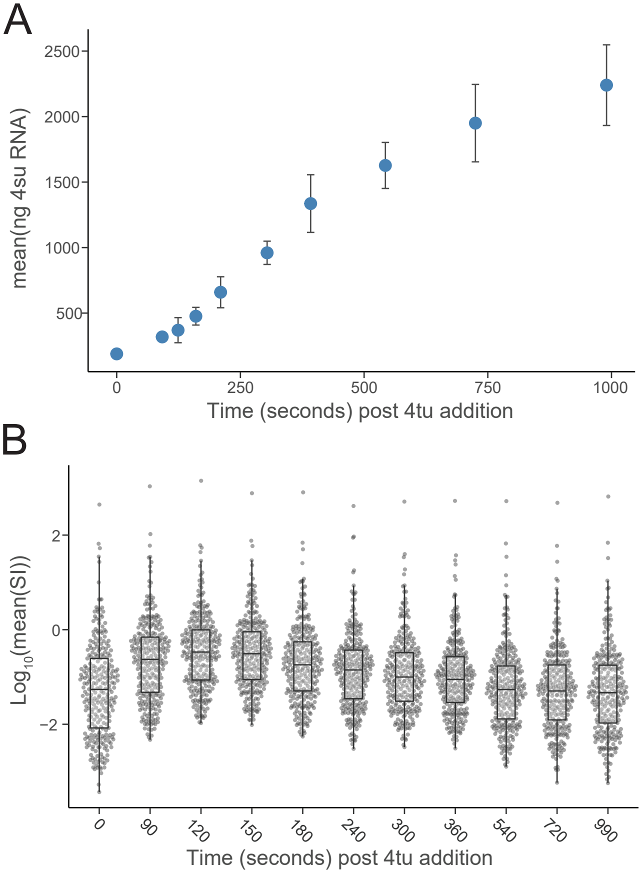

### Figure S2

A

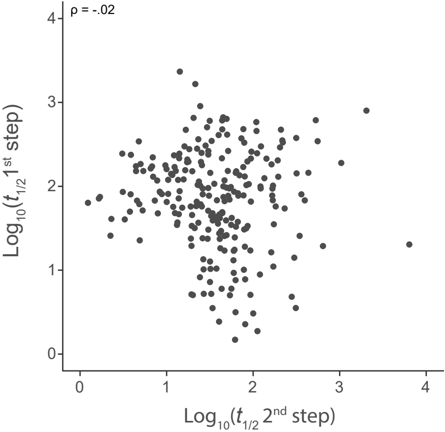

### Figure S3

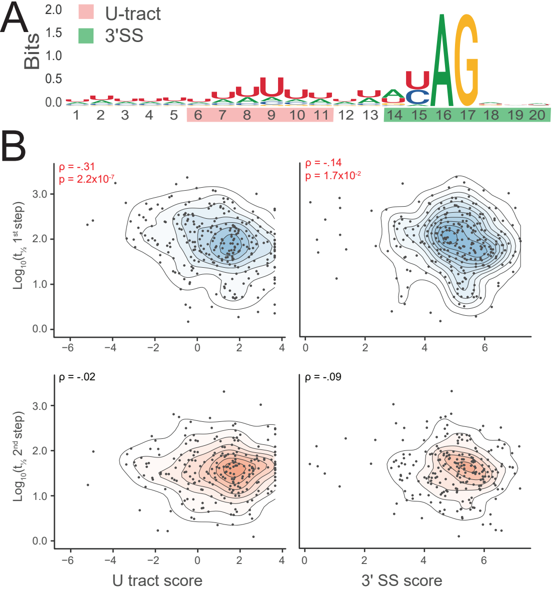

### Figure S4

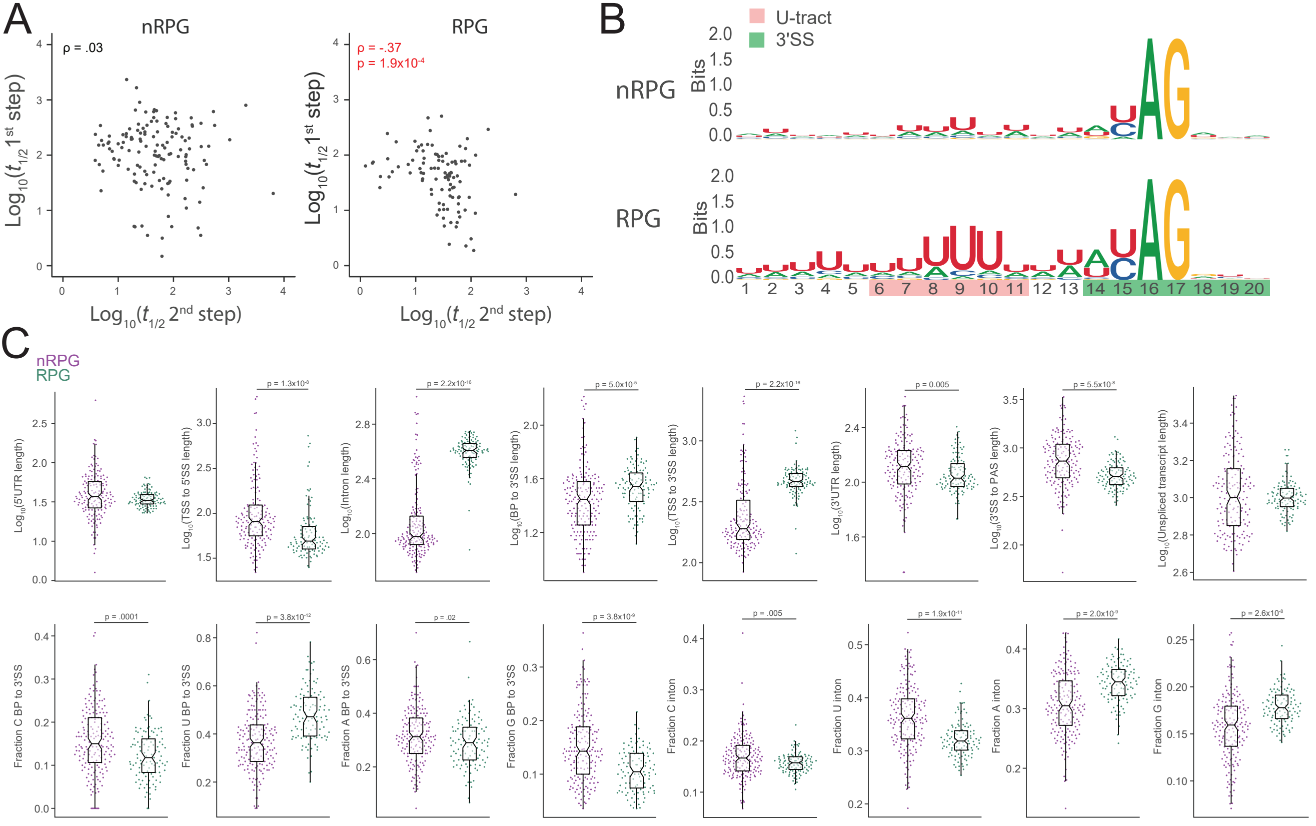
